# Liming enhances the abundance and stability of nitrogen-cycling microbes: The buffering effect of long-term lime application

**DOI:** 10.1101/2024.04.21.590279

**Authors:** Akari Mitsuta, Kesia Silva Lourenco, Jingjing Chang, Mart Ros, René Schils, Yoshitaka Uchida, Eiko Eurya Kuramae

## Abstract

Lime application (liming) has historically been used to ameliorate soil acidity in grasslands. Liming effectively improves soil pH, plant productivity, and soil physicochemical properties, but the long-term impact of acidity control by liming on key microbial nitrogen (N)-cycling genes in semi-natural grasslands is unknown. We investigated the effect of 65 years of liming on N-cycling processes in the limed and control plots of the Ossekampen long-term grassland experiment in the Netherlands. These plots have not received any other fertilizers for 65 years. Soil sampling and nitrous oxide (N_2_O) emission measurements were conducted three times in spring and four times in summer, and quantitative real-time PCR was performed to determine the abundances of N-cycling genes, including ammonia-oxidation (*amoA*), denitrification (*nirS*, *nirK*, *nosZ*), and N-fixation (*nifH*) genes. Long-term liming increased the abundances of nitrifiers and denitrifiers but did not increase N_2_O emissions. Additionally, liming had a buffering effect that stabilized the population of N-cycling microbes against seasonal variations in abundance. Our results indicate that improving soil acidity through liming facilitates microbial N-cycling processes without increasing N_2_O emissions.

**Highlights:**

- 65 years of liming increased bacterial but not archaeal and fungal community abundance.
- Liming increased the abundance of microbial N-cycling genes.
- The buffering effect of liming reduced seasonal variations in the abundance of N microbes.

## 1. Introduction

Approximately 40% of the world’s arable land is occupied by acidic soil, usually defined as a soil pH below 5.5 (Bian et al., 2013; Uexküll and Mutert, 1995). Acidic soil is associated with lower macronutrient availability, plant production, and soil microbial diversity and abundance (Bossolani et al., 2021, 2020; Rousk et al., 2010; Shen et al., 2013). In grasslands, the negative effects of soil acidity have historically been ameliorated by applying lime (liming). Lime neutralizes soil acidity by producing CO_2_ and H_2_O, effectively raising the soil pH and improving soil physical properties, nutrient availability, and plant productivity (Abdalla et al., 2022; Bossolani et al., 2020; Haynes and Naidu, 1998). Liming can also improve microbial abundance and community diversity (Jiang et al., 2021; Kennedy et al., 2004; Xue et al., 2010). Furthermore, in agricultural soil, acidity is associated with increased emissions of nitrous oxide (N_2_O) (Shaaban et al., 2019; Weier and Gilliam, 1986; Yamulki et al., 1997), a gas with a global warming potential 273 times greater than that of carbon dioxide (CO_2_) and a lifetime of approximately 120 years (Forster et al., 2021). Regardless of the application of nitrogen (N) fertilizer, liming is estimated to reduce N_2_O emissions from acidic agricultural soil by 26% by enhancing the activity of microbes that reduce N_2_O to N_2_ (Wang et al., 2021).

Because N is an essential, growth-limiting nutrient for plant production, studies of the effect of liming on microbial N-cycling processes are important. Liming affects soil inorganic N content and gaseous losses of N_2_O by influencing the abundances of nitrifying and denitrifying microbes. The first rate-limiting step in nitrification is ammonia oxidation, which is catalyzed by the enzyme ammonia monooxygenase in ammonia-oxidizing bacteria (AOB), ammonia-oxidizing archaea (AOA), and complete ammonia-oxidizing (comammox) bacteria (Daims et al., 2015; Prosser and Nicol, 2008; Purkhold et al., 2000). Soil pH affects the community composition and activity of AOB and AOA (Nicol et al., 2008): AOB often predominate under conditions of high N availability and neutral soil pH, whereas AOA are more prevalent under conditions of lower N availability and acidic soil. Several studies have reported increases in AOB abundance in response to rising soil pH due to liming (Teutscherova et al., 2017; Wakelin et al., 2009; Yao et al., 2011). Liming also increases the abundance of comammox bacteria (Wang et al., 2023), although information on these recently discovered bacteria is limited (Daims et al., 2015). In addition to affecting nitrification, liming influences denitrification, a process that converts nitrate to nitrogen gas (N_2_) while producing N_2_O. The fraction of incomplete denitrification, which results in N_2_O production, is often higher in acidic soils than in neutral soils (Čuhel et al., 2010; Dannenmann et al., 2008; Liu et al., 2010), primarily due to the inhibitory effect of soil acidity on the enzyme N_2_O reductase (Frostegård et al., 2022). Thus, improving soil pH through liming reportedly decreases N_2_O emissions by promoting complete denitrification (Wang et al., 2021; Zhang et al., 2022). However, most studies of the effects of liming have been conducted over the short-term, generally less than 30 years. A comparison of the long- and short-term impacts of liming found that application duration is an important factor controlling denitrification and N_2_O emissions, likely due to adaptation of the microbial community (Baggs et al., 2010). Thus, further study is needed to assess the long-term effects of liming on key microbes involved in N-cycling processes.

In addition to influencing microbial nitrifiers and denitrifiers, liming affects the abundance and activity of microorganisms performing biological N fixation (diazotrophs). Several field studies have reported positive effects of liming on diazotroph abundance in grasslands, rotation croplands, and rice paddy fields (Bossolani et al., 2020; Jiang et al., 2022; Wakelin et al., 2009). However, liming has also been shown to reduce *nifH* gene abundance (Lin et al., 2018; Silveira et al., 2021), suggesting that our understanding of the effects of liming on diazotrophs is not consolidated. These discrepancies may be attributable to differences in experimental setup and duration, as the succession and stabilization of microbial community structures are lengthy processes (Geisseler and Scow, 2014). Thus, long-term experiments may provide valuable insights into microbial community dynamics and foster consensus on the effects of liming.

The soil microbial community and functional microbial community involved in N cycling in agricultural ecosystems may also undergo seasonal changes (Kennedy et al., 2005; Madegwa and Uchida, 2021a). Soil microbial community structure adjusts with changing seasons, primarily due to shifts in soil physicochemical characteristics such as temperature, moisture, and nutrient availability. Because liming enhances soil physicochemical stability, including soil pH, nutrient availability, soil moisture retention (Chan and Heenan, 1999; Hati et al., 2008), and soil aggregation (Blomquist et al., 2022; Keiblinger et al., 2016), it may contribute to increased stability within microbial functional groups. In fact, Madegwa and Uchida (2021b) demonstrated that liming enhances the stability of the soil microbial community in response to disturbances caused by organic and chemical fertilizer applications. However, few studies have investigated the seasonal impact of liming, and the contribution of long-term liming to seasonal variations in microbial N gene abundance remains an open question.

In the present study, we addressed these gaps in knowledge by evaluating the effects of long-term (65 years) liming on the abundances of microbial genes involved in N cycling, including ammonia oxidation (*amoA*), denitrification (*nirS*, *nirK*), and N fixation (*nifH*), as determined by quantitative real-time PCR (qPCR). We hypothesized that (i) long-term liming increases the abundances of microbial N-cycle genes related to nitrification, denitrification, and N fixation; (ii) liming mitigates seasonal effects on the abundances of microbial N-cycle genes; and (iii) liming decreases N_2_O emissions by increasing the abundance of complete N_2_O reductase (*nosZ*) genes.

## 2. Materials and Methods

### 2.1 Site description

The Ossekampen long-term grassland experiment (51 degrees 58′15″N; 5 degrees 38′18″E, Wageningen, The Netherlands) was started in 1958 in an extensively grazed species-rich grassland with soil characterized as heavy river clay. Prior to the experiment, the land was grazed and had been used in alternate years for haymaking. The treatments consist of lime (grounded calcium carbonate, 357 kg Ca ha^−1^ yr^−1^) and control (no fertilizer application) in duplicate 40-m^2^ (16 m× 2.5 m) plots. The grass is mown twice yearly, in July and October, and the biomass is removed from a 2.5-m-wide perimeter to prevent the dispersal of seeds into other plots. The plots are separated by unfertilized 2.5-m-wide buffer strips to avoid contamination between treatments, and these strips are similarly maintained by mowing.

### 2.2 Soil sampling and greenhouse gas measurement

Soil samples (150 g) were collected from the 0–10 cm top layer in 2022. For both soil and gas sampling, each plot was split into two sub-plots and duplicated. Sampling was conducted three times in spring (April 6, 13, and 19) and four times in summer (July 7, 14, and 18 and August 2). Thus, a total of 56 samples (2 treatments × 4 replications × 7 time points) were collected. Immediately after sampling, soil subsamples (30 g) were stored at −80°C until the molecular analyses, and the remaining samples were stored at −20°C for use in chemical property analyses. Lime was applied on April 11.

In parallel with soil sampling, CO_2_ and N_2_O fluxes were measured with a PICARRO G2508 ring-down spectroscopy gas analyzer (Picarro Inc., Santa Clara, CA, USA). These fluxes were measured with a closed chamber technique. One week before the measurement, polyvinylchloride (PVC) rings with an inner diameter of 19 cm and a height of 10 cm were inserted 5 cm into the soil, 1.5 m from both edges of each plot. For the flux measurements, a white opaque polypropylene chamber with a diameter of 20 cm and a height of 11 cm was fastened on the PVC ring to create a flux chamber. The flux was determined assuming a linear increase over the incubation period. The validity of this assumption was checked by monitoring the concentration changes of one or two measuring points continuously over the incubation time.

### 2.3 Soil chemical property analysis

Soil mineral N (NH_4_⁺₋N, NO_2_ ^−^₋N, and NO_3_ ^−^₋N) was measured with a continuous flow analytical system (Skalar San^++^ continuous flow meter) after extraction with 2 M KCl in 1:5 soil:solution. Soil pH was measured in 0.01 M CaCl_2_ due to the advantage of damping the error caused by seasonal variations in soil salt concentration (Kome et al., 2018). Available P-phosphate was determined by calorimetry, and exchangeable cations (K^+^, Ca^2+^, and Mg^+^) were determined by atomic absorption spectrometry (Shimadzu AA-7000) analysis of the soil extract obtained using ion-exchange resin (van Raij et al., 2001). Cationic micronutrients [iron (Fe), manganese (Mn), copper (Cu) and zinc (Zn)] were extracted in a solution (pH 7.3) containing 0.005 M diethylenetriaminepentaacetic acid (DTPA), 0.1 M triethanolamine (TEA), and 0.01 M CaCl_2_ and determined by atomic absorption spectrometry.

### 2.4 DNA extraction and quantification of N-cycle gene abundance

Total soil DNA was extracted from 0.40 g of soil using the MoBio PowerSoil DNA Isolation Kit (MoBio, Solana Beach, CA, USA) according to the manufacturer’s instructions. DNA quantity and quality were determined using a NanoDrop ND-1000 spectrophotometer (NanoDrop Technologies, DE, USA).

The abundances of the functional genes *amoA*-AOB, *amoA*-AOA, *amoA*-comammox, *nirS* clusters I-IV, *nirK* clusters I-IV, *nosZ* clades I and II, and *nifH* and ribosomal RNA genes for total bacteria, archaea, and fungi were quantified by qPCR using the CFX96 Touch™ Real-Time PCR Detection System (Bio-Rad). The qPCR standard curve of each functional gene was constructed as described in the supplemental material. For each primer set, the cycling conditions, annealing temperature, cycle number, and primer amount were optimized, and the amplicon of the target gene was checked with electrophoresis (Table S1). The genes *nirS* clusters II-IV and *nirK* clusters III and IV were not detected in our samples.

### 2.5 Statistical analysis

All statistical analyses were conducted in R version 4.3.0. A linear mixed model was used to determine the significance of differences in microbial gene abundance and chemical properties between treatments (control and lime) and seasons (spring and summer). The model was calculated with the “lme4” package (Bates et al., 2015) with treatment and season as fixed effects and time and site as random effects. When the interaction effect of fertilizer and season was significant (*P* < 0.05), pairwise comparisons were conducted with the “emmeans” package (Lenth et al., 2020). Redundancy analysis (RDA) was performed to identify chemical properties that significantly affected the abundance of microbial genes in the “vegan” package (Dixon, 2003). The microbial genes and chemical properties listed in Tables 1 and 2 were included in the analysis. Permutational multivariate analysis of variance [PERMANOVA; Anderson (2001)] was performed to determine whether the factors of treatment and season influenced microbial gene abundances. In addition, homogeneity of multivariate dispersion tests [PERMDISP; Anderson et al (2006)] were conducted to evaluate differences in dispersion between treatment types. Both PERMANOVA and PERMDISP were conducted using Bray–Curtis distance metrics with 10,000 permutations. Correlations between the abundance of each gene and chemical properties were tested with Spearman’s coefficient analysis. Furthermore, to determine which genes were most impacted by liming, a volcano plot that expressed significant differences in gene abundance as the fold change between treatment types was constructed.

**Table 1.** Microbial gene abundances in each treatment and season (mean of log-transformed gene copy number ± standard deviation).

| Gene | ANOVA |  |  | Control |  | Lime |  |
| --- | --- | --- | --- | --- | --- | --- | --- |
| | Treatment | Season | Treatment×season | Spring | Summer | Spring | Summer |
| Archaeal 16S rRNA | ns | ** | * | 7.8 ± 0.032a | 7.5 ± 0.038b | 7.9 ± 0.032a | 7.6 ± 0.026b |
| Bacterial 16S rRNA | * | * | ** | 9.2 ± 0.030a | 9.1 ± 0.025b | 9.3 ± 0.018a | 9.2 ± 0.017a |
| Fungal 18S rRNA | * | ns | ns | 8.3 ± 0.038 | 8.4 ± 0.035 | 8.1 ± 0.038 | 8.2 ± 0.035 |
| *amoA*-AOB | *** | ns | ns | 3.0 ± 0.11 | 3.3 ± 0.20 | 4.6 ± 0.123 | 4.7 ± 0.061 |
| *amoA*-AOA | ns | ns | ns | 5.1 ± 0.23 | 4.7 ± 0.13 | 5.6 ± 0.14 | 5.4 ± 0.13 |
| *amoA*-comammox | *** | * | ns | 4.2 ± 0.056 | 3.9 ± 0.079 | 5.4 ± 0.118 | 5.2 ± 0.088 |
| *nifH* | ns | * | ** | 7.8 ± 0.080a | 7.5 ± 0.063b | 7.8 ± 0.071a | 7.8 ± 0.039a |
| *nirK* Cluster I | ns | * | ns | 7.2 ± 0.043 | 7.1 ± 0.055 | 7.2 ± 0.029 | 7.1 ± 0.026 |
| *nirK* Cluster II | *** | * | ns | 7.4 ± 0.071 | 7.1 ± 0.036 | 7.9 ± 0.019 | 7.7 ± 0.026 |
| *nirS* Cluster I | ** | *** | ns | 7.8 ± 0.042 | 7.7 ± 0.039 | 8.1 ± 0.048 | 8.1 ± 0.026 |
| *nosZ* Clade I | ns | ** | ns | 7.0 ± 0.056 | 6.6 ± 0.052 | 7.1 ± 0.037 | 6.7 ± 0.035 |
| *nosZ* Clade II | ** | ns | ns | 6.3 ± 0.031 | 6.3 ± 0.045 | 6.6 ± 0.034 | 6.6 ± 0.034 |

The asterisk indicates the *P* value for the linear mixed model with factors of treatment and season (\**P* < 0.05, \*\**P* < 0.01, and \*\*\**P* < 0.001; “ns” indicates not significant). Values for the same gene and treatment with different lowercase letters are significantly different (*P* < 0.05) between spring and summer.

## 3. Results

### 3.1 Abundance of N-cycle functional genes

The abundances of the microbial *amoA-*AOB, *amoA-*comammox, *nirK* cluster II, *nirS* cluster I, and *nosZ* clade II genes were significantly higher in the liming treatment than in the control in both spring and summer (Table 1). The abundance of the *nifH* gene was higher in the liming treatment than in the control in summer but did not differ between treatments in spring. In both seasons, fungal abundance was significantly lower in the liming treatment than in the control.

The volcano plot in Figure 1 provides a visual representation of the differences in gene abundance between the liming treatment and the control. Points located outside the center (0) indicate significant differences in response to liming. Genes that are positioned further to the right exhibited a stronger positive response to liming, and genes that are positioned closer to the top were stably more abundant in the liming treatment over the sampling period. In summary, liming had greater positive effects on the abundances of *amoA*-AOB, *amoA*-comammox, and *nirK* cluster II than on the other genes.

**Fig. 1.**
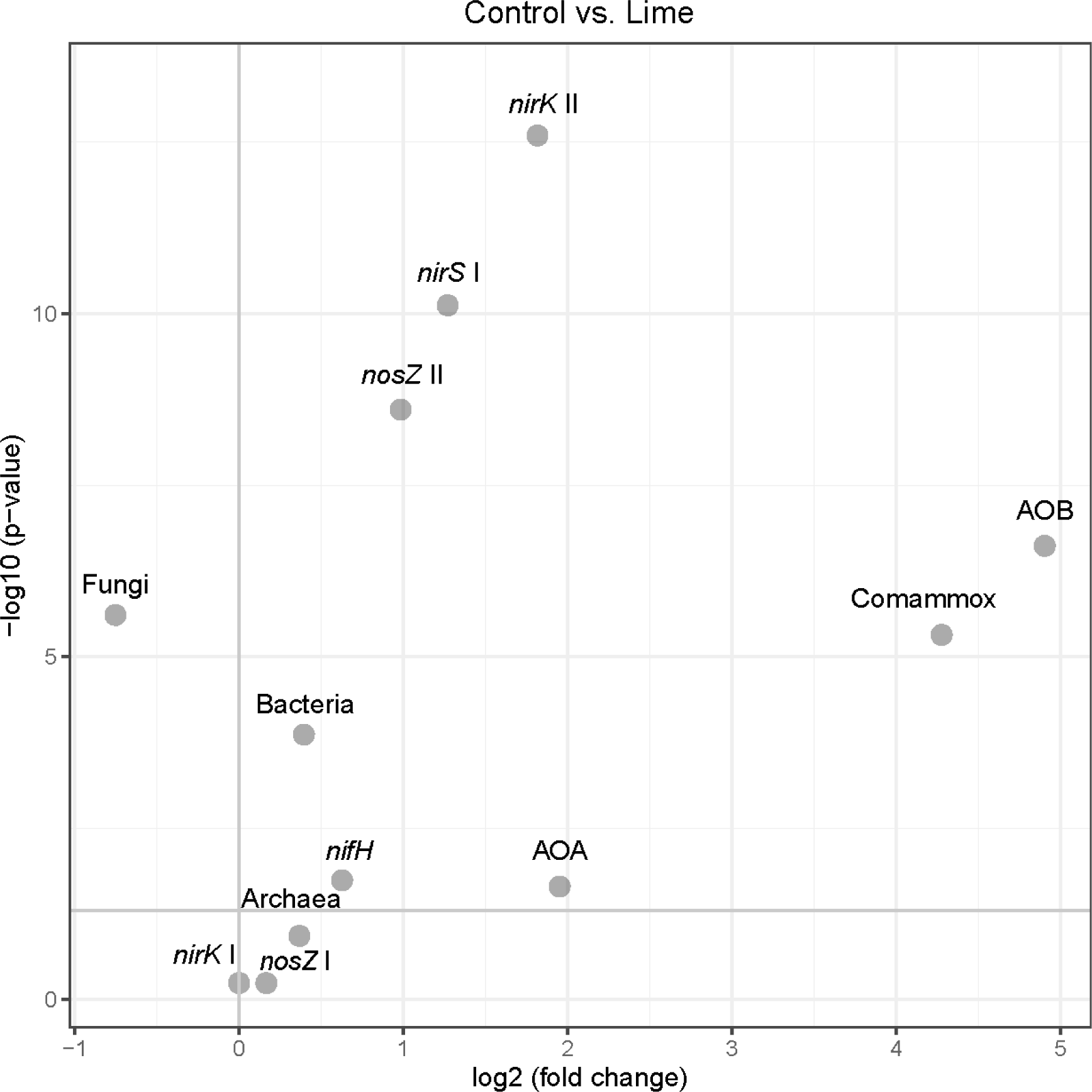
Volcano plot of differences in microbial gene abundance between the control and lime treatments. The y axis represents the negative log10 of the *P* value calculated with the linear mixed model, and the x axis represents the log of the fold change between the treatments.

### 3.2 RDA plot and stability of microbial gene abundance

The RDA plot in Figure 2a indicates the relationship between soil chemical properties and the relative abundance of microbial genes. According to PERMANOVA, gene abundances differed significantly (*P* < 0.001) between treatments and seasons. However, no interaction between treatment and season was detected. To evaluate differences in the dispersion of gene abundances, PERMDISP analysis was conducted. A significant (*P* < 0.001) effect of treatment was observed, indicating that the seasonal dispersion of gene abundance was lower in the liming treatment than in the control. This suggests that liming had a buffering effect, i.e., reduced the impact of seasonal variations on gene abundance in the soil. According to the RDA, the differences in microbial gene abundance between the liming and control treatments could be explained by soil pH, moisture, and nutrients, i.e., K^+^, Ca^2+^, Mg^2+^, Cu, Fe, Mn, and Zn (Fig. 2b).

**Fig. 2.**
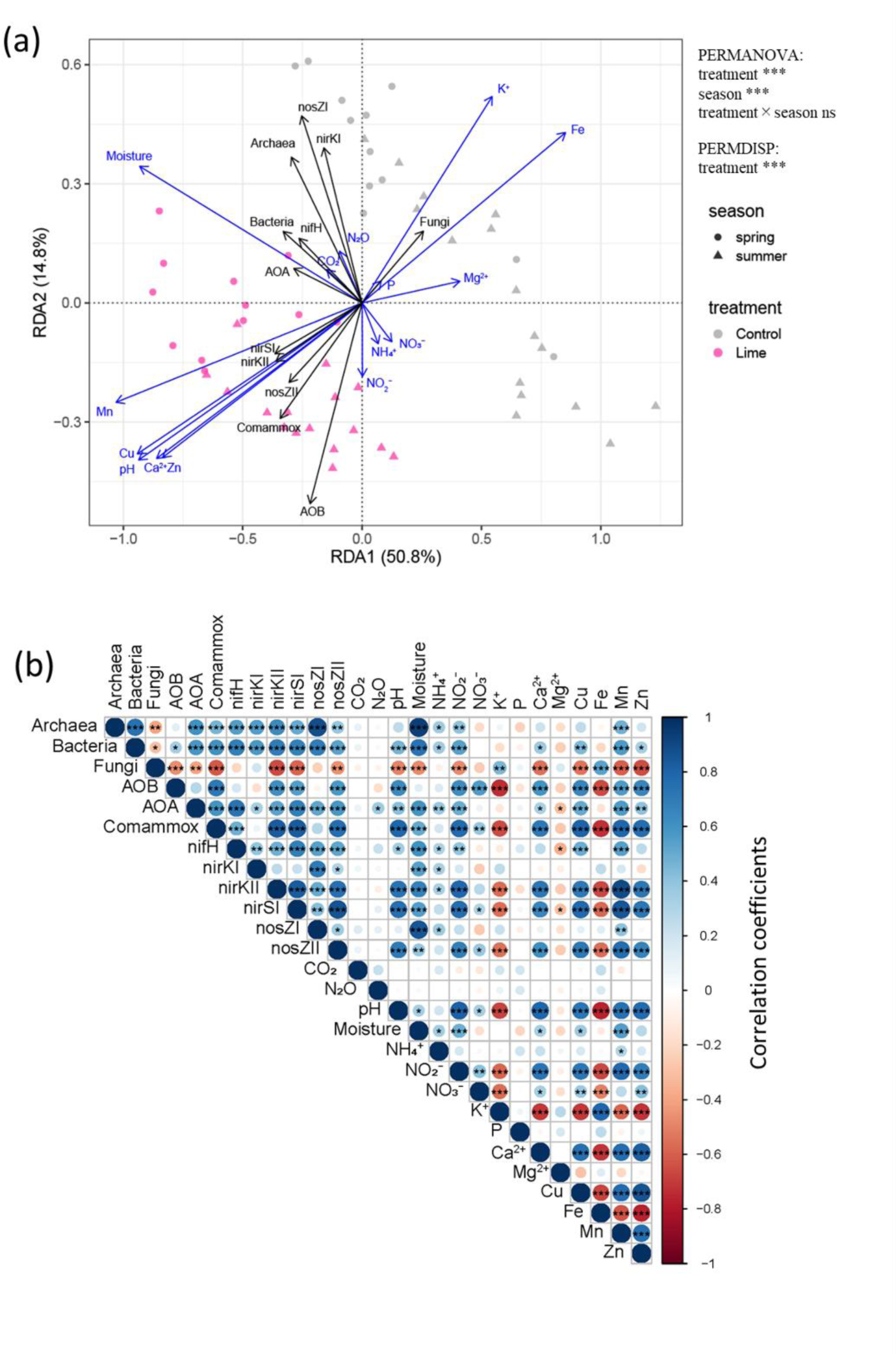
(a) Redundancy analysis (RDA) of the relative abundances of N-cycle functional genes and soil chemical properties. Only significant vectors (*P* < 0.05) for chemical properties are shown with blue arrows. Significant clustering by treatment and season identified by PERMANOVA and significant differences in sample dispersion between treatments identified by PERMDISP are shown in the upper right corner (\*\*\**P* < 0.001). (b) Heatmap of Spearman’s correlation coefficients between microbial gene abundances, soil chemical properties, and greenhouse gas emissions with *P* values (\**P* < 0.05, \*\**P* < 0.01, \*\*\**P* < 0.001).

### 3.3 Soil chemical properties, greenhouse gas emissions, and plant production

As expected, soil pH was significantly (*P* < 0.001) higher in the liming treatment than in the control (Table 2). Soil moisture content was also significantly (*P* < 0.05) higher in the liming treatment than in the control. The soil concentrations of NH_4_⁺₋N did not differ between the treatments but NO_2_ ^−^₋N, and NO_3_ ^−^₋N were significantly (*P* < 0.001) higher in the liming treatment than in the control. Compared with the control, soil Ca^2+^, Cu, Mn, and Zn concentrations were significantly higher in the liming treatment, whereas soil K^+^ and Fe concentrations were significantly lower in the liming treatment.

**Table 2.**
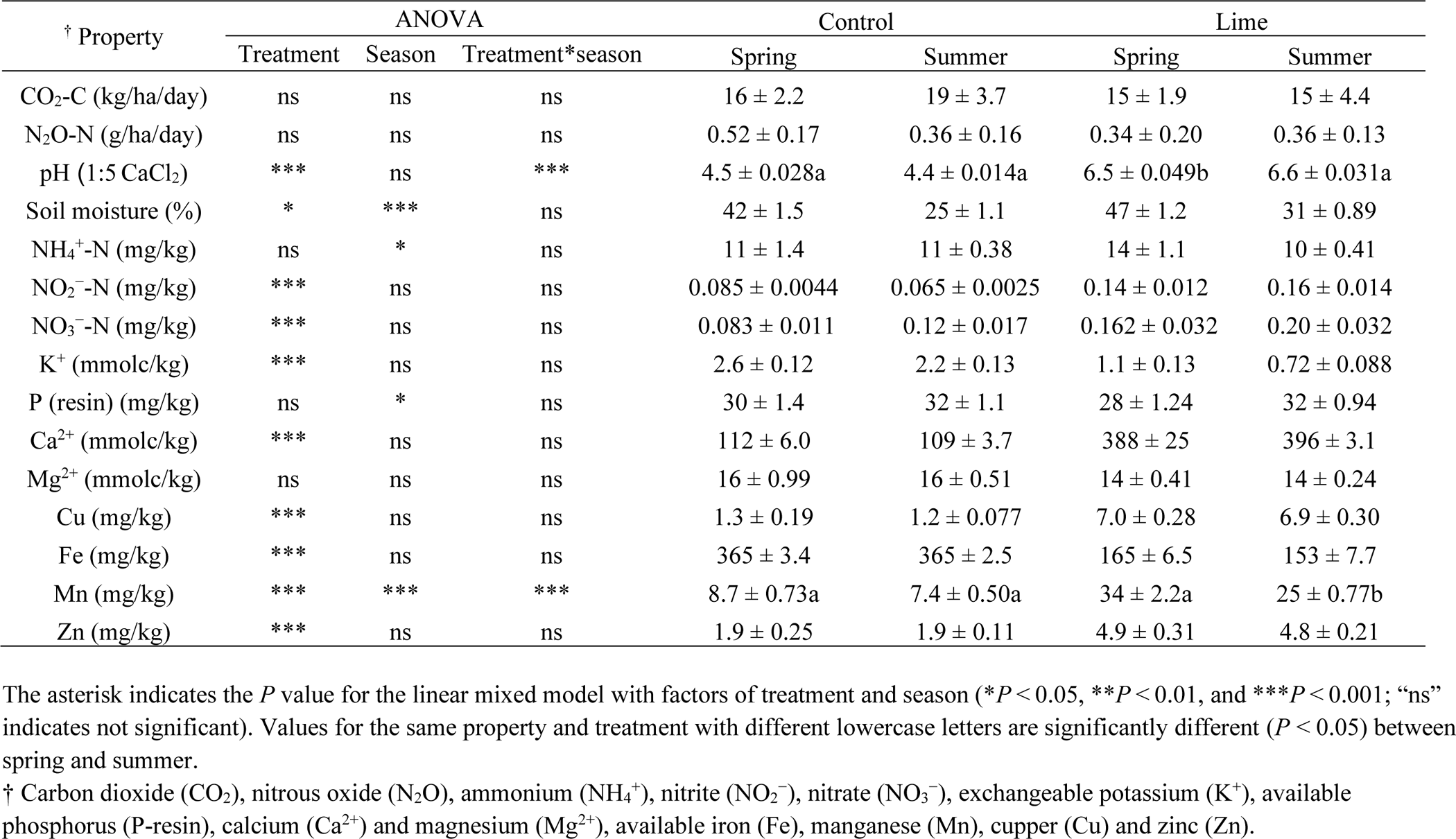
Effect of lime application on greenhouse gas (GHG) emissions and soil chemical properties (mean ± standard deviation).

| † Property | ANOVA |  |  | Control |  | Lime |  |
| --- | --- | --- | --- | --- | --- | --- | --- |
|  | Treatment | Season | Treatment*season | Spring | Summer | Spring | Summer |
| CO<sub>2</sub>-C (kg/ha/day) | ns | ns | ns | 16 ± 2.2 | 19 ± 3.7 | 15 ± 1.9 | 15 ± 4.4 |
| N<sub>2</sub>O-N (g/ha/day) | ns | ns | ns | 0.52 ± 0.17 | 0.36 ± 0.16 | 0.34 ± 0.20 | 0.36 ± 0.13 |
| pH (1:5 CaCl<sub>2</sub>) | *** | ns | *** | 4.5 ± 0.028a | 4.4 ± 0.014a | 6.5 ± 0.049b | 6.6 ± 0.031a |
| Soil moisture (%) | * | *** | ns | 42 ± 1.5 | 25 ± 1.1 | 47 ± 1.2 | 31 ± 0.89 |
| NH<sub>4</sub><sup>+</sup>-N (mg/kg) | ns | * | ns | 11 ± 1.4 | 11 ± 0.38 | 14 ± 1.1 | 10 ± 0.41 |
| NO<sub>2</sub><sup>-</sup>-N (mg/kg) | *** | ns | ns | 0.085 ± 0.0044 | 0.065 ± 0.0025 | 0.14 ± 0.012 | 0.16 ± 0.014 |
| NO<sub>3</sub><sup>-</sup>-N (mg/kg) | *** | ns | ns | 0.083 ± 0.011 | 0.12 ± 0.017 | 0.162 ± 0.032 | 0.20 ± 0.032 |
| K<sup>+</sup> (mmolc/kg) | *** | ns | ns | 2.6 ± 0.12 | 2.2 ± 0.13 | 1.1 ± 0.13 | 0.72 ± 0.088 |
| P (resin) (mg/kg) | ns | * | ns | 30 ± 1.4 | 32 ± 1.1 | 28 ± 1.24 | 32 ± 0.94 |
| Ca<sup>2+</sup> (mmolc/kg) | *** | ns | ns | 112 ± 6.0 | 109 ± 3.7 | 388 ± 25 | 396 ± 3.1 |
| Mg<sup>2+</sup> (mmolc/kg) | ns | ns | ns | 16 ± 0.99 | 16 ± 0.51 | 14 ± 0.41 | 14 ± 0.24 |
| Cu (mg/kg) | *** | ns | ns | 1.3 ± 0.19 | 1.2 ± 0.077 | 7.0 ± 0.28 | 6.9 ± 0.30 |
| Fe (mg/kg) | *** | ns | ns | 365 ± 3.4 | 365 ± 2.5 | 165 ± 6.5 | 153 ± 7.7 |
| Mn (mg/kg) | *** | *** | *** | 8.7 ± 0.73a | 7.4 ± 0.50a | 34 ± 2.2a | 25 ± 0.77b |
| Zn (mg/kg) | *** | ns | ns | 1.9 ± 0.25 | 1.9 ± 0.11 | 4.9 ± 0.31 | 4.8 ± 0.21 |

The asterisk indicates the *P* value for the linear mixed model with factors of treatment and season (\**P* < 0.05, \*\**P* < 0.01, and \*\*\**P* < 0.001; “ns” indicates not significant). Values for the same property and treatment with different lowercase letters are significantly different (*P* < 0.05) between spring and summer.
† Carbon dioxide (CO<sub>2</sub>), nitrous oxide (N<sub>2</sub>O), ammonium (NH<sub>4</sub><sup>+</sup>), nitrite (NO<sub>2</sub><sup>-</sup>), nitrate (NO<sub>3</sub><sup>-</sup>), exchangeable potassium (K<sup>+</sup>), available phosphorus (P-resin), calcium (Ca<sup>2+</sup>) and magnesium (Mg<sup>2+</sup>), available iron (Fe), manganese (Mn), copper (Cu) and zinc (Zn).

Soil pH was positively correlated with the abundances of *amoA*-AOB, *amoA*-AOA, *amoA*-comammox, *nifH*, *nirK* cluster II, and *nirS* cluster I (Fig. 2b). Soil moisture was positively correlated with the abundances of *amoA*-AOA, *amoA*-comammox, *nifH*, *nirK* clusters I and II, *nirS* cluster I, and *nosZ* clades I and II. Among soil nutrients, the concentrations of Ca^2+^, Cu, Mn, and Zn were significantly higher in the liming treatment than in the control and were positively correlated with the abundances of *amoA*-AOB, *amoA*-AOA, *amoA*-comammox, *nirK* cluster II, *nirS* cluster I, and *nosZ* clade II. Additionally, the soil concentrations of Cu and Mn were positively correlated with *nifH* gene abundance.

CO_2_ and N_2_O emissions, which were measured in parallel with soil sampling, were not significantly different between the liming and control treatments during the experimental period.

After mowing in July and October, grasses were harvested, and biomass production was determined (Fig. 3). Compared with the control, the aboveground dry matter yield in the liming treatments was 52% higher in July and 154% higher in October. Moreover, the N yield from the plants was 61.6% higher in July and 193% higher in October in the liming treatment than in the control.

**Fig. 3.**
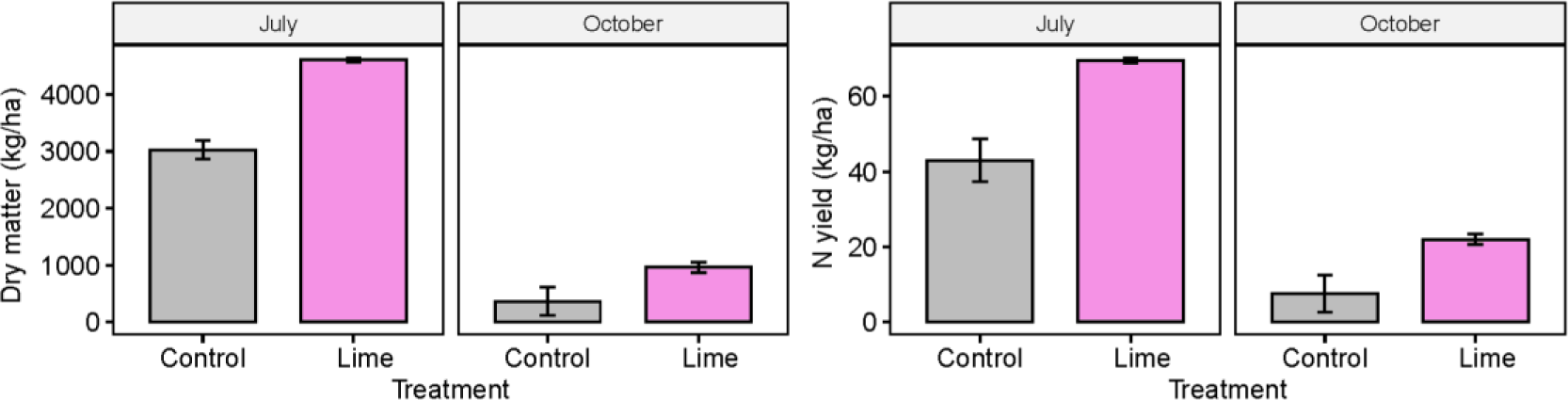
Dry matter and nitrogen yields of plants measured at grass mowing in July and October.

## 4. Discussion

### 4.1 Effect of liming on microbial N-cycle gene abundance

We found that long-term lime application enhanced the abundances of microbial genes involved in nitrification (*amoA*-AOB and *amoA*-comammox) and denitrification (*nirK* cluster II, *nirS* cluster I, and *nosZ* clade II). These results are consistent with previous observations that liming enhances the abundances of nitrifiers (Teutscherova et al., 2017; Wakelin et al., 2009) and denitrifiers (Jha et al., 2020; Vázquez et al., 2020), however few studies have investigated the effect of liming on comammox bacteria. We found a strong positive impact of liming on comammox gene abundance, emphasizing the need for further exploration of the responses of comammox bacteria to liming. The positive effect of liming on the abundances of microbial N-cycle genes can be attributed to increased soil pH, which is a pivotal determinant of various aspects of soil biogeochemical cycles, such as micronutrient availability and soil organic matter mineralization (Kemmitt et al., 2006; Silveira et al., 2021). Additionally, lime helps improve soil aggregation and maintain soil moisture, which are important for microbial activity (Chan and Heenan, 1999; Hati et al., 2008). Our results suggest that 65 years of long-term liming had substantial positive effects on soil pH (increasing from 4.5 to 6.6), soil micronutrient availability (particularly increases in the availability of Ca^2+^, Mn, and Zn), and soil moisture content. As a result, liming enhanced the abundances of microbial genes involved in the N cycle.

Furthermore, liming significantly mitigated the reduction in *nifH* gene abundance in summer (Table 1). We observed significant positive correlations of *nifH* gene abundance with soil moisture and Cu and Mn contents. These correlations can be explained by the properties of nitrogenase, the oxygen-labile enzyme encoded by *nifH* that is responsible for biological N fixation. Low soil moisture, especially moisture content below the soil’s water-holding capacity, can hinder biological N fixation (Burris and Roberts, 1993; Groß et al., 2022; Smercina et al., 2019). Cu and Mn play crucial roles in protecting diazotrophic proteins from oxidation by acting as cofactors for the enzyme superoxide dismutase, which catalyzes the dismutation of superoxide radicals to oxygen (Rubio et al., 2004). Thus, the positive effects of liming on soil moisture, Cu, and Mn were essential for maintaining *nifH* gene abundance during the dry summer season. Our findings differ from those of Bossolani et al. (2020), who found positive relationships of *nifH* gene abundance with exchangeable Ca^2+^ and Mg^2+^ contents but no relationships with Cu and Mn contents. However, in the present study, the concentrations of Cu and Mn in the control treatment were lower (1.3 and 8.7 mg/kg, respectively) than those in previous studies (Bossolani et al., 2020; Brennan et al., 2008; Min et al., 2022) in which Fe dominated due to the strong acidity of the soil (Truog, 1947). These differences are attributable to variations in soil types; for example, the study by Bossolani et al. (2020) was conducted in tropical agricultural soil, whereas this study was conducted in a temperate semi-natural grassland. Our findings indicate that soil type should be considered when examining the impact of liming on diazotroph abundance.

### 4.2 Effect of liming on seasonal variations in the abundance of N microbes

Under liming, seasonal variations in soil microbial gene abundances were reduced, indicating that liming buffered microbial gene abundance against seasonal fluctuations (Fig. 2). The significant correlations between most of the functional genes and soil moisture content imply that the buffering effect of lime on soil moisture was a key factor directly or indirectly affecting the stability of gene abundance. Compared with spring, soil moisture decreased significantly in summer in both the liming treatment (from 47% to 31%) and the control (from 42% to 25%). However, the reduction in soil moisture was less pronounced in the liming treatment. Similar effects of liming on the entire microbial community have been reported; specifically, liming reduced the changes in microbial community structure caused by the application of organic and inorganic fertilizers (Madegwa and Uchida, 2021b). Given these results and our observations, we suggest that liming stabilizes soil microorganism abundance by buffering variations in the environment caused by fertilizer application and seasonal changes. Further research is crucial to unravel how liming stabilizes microbial functions related to other cycles, i.e., P, C and S, or microbial communities that influence nutrient cycles.

### 4.3 Effect of liming on N_2_O emissions

Interestingly, N_2_O emissions did not differ significantly between the liming and control treatments, despite the increased abundances of genes related to nitrification and denitrification in the liming treatment. The lack of increase in N_2_O emissions under liming may be attributable to the increased abundance of *nosZ* clade II, which encodes the N_2_O reductase enzyme. Microbes with *nosZ* clade II genes typically lack genes related to N_2_O production and are primarily associated with N_2_O consumption (Jones et al., 2014; Shan et al., 2021). By contrast, the abundance of *nosZ* clade I genes did not differ significantly between treatments. Instead, the primary influencing factor was seasonal variations, as *nosZ* clade I gene abundance was positively correlated with soil moisture content. These findings are in agreement with previous results demonstrating that *nosZ* clade II abundance is more strongly influenced by environmental factors such as soil pH (Domeignoz-Hortal et al., 2015; Jones et al., 2014), whereas *nosZ* clade I abundance is more strongly influenced by soil moisture than soil chemical properties (Xu et al., 2020). Our results support these findings and further contribute to our understanding of niche partitioning within *nosZ* clades. However, it should be noted that mineral N fertilizer was not applied in the plots in our study; thus, available N was low.

Additionally, we observed reduced fungal abundance in the liming treatment. Fungi contribute to N_2_O emissions (Aldossari and Ishii, 2021), and the ratio of fungi to bacteria is usually higher in acidic soils, potentially leading to increased fungus-mediated N_2_O emissions (Chen et al., 2015; Huang et al., 2021; Lourenço et al., 2018; Rütting et al., 2013). The reduced fungal abundance in the liming treatment may have contributed to the lack of a significant difference in N_2_O emissions despite the increased abundances of nitrifiers and denitrifiers. Moreover, the role of nitrifier denitrification, which could be a primary pathway for N_2_O production in lime-treated soil, should be considered. Several studies have linked increased N_2_O emissions in lime-treated soil to increased nitrifier denitrification, especially under high N availability, e.g., urine application at 500 kg N/ha (Clough et al., 2003; Khan et al., 2011). Although we observed a significant increase in *amoA*-AOB gene abundance in the liming treatment, our results suggest that the contribution of nitrifier denitrification to N_2_O production was limited by low N availability. Overall, the influence of liming on N_2_O emissions depends on the net effect on N_2_O reductases, fungal denitrification, and nitrifier denitrification.

Our study demonstrates that long-term liming is advantageous for sustaining the N cycle in acidic grasslands. This study is unique in that it evaluates N-cycle processes from fixation to emission following a very long period of liming—65 years—in an acidic semi-natural grassland. Earlier studies of the same grassland indicated that liming increased plant and bacterial diversity (Cassman et al., 2016), and our study provides strong evidence that 65 years of liming has greatly shaped this grassland ecosystem by maintaining a diverse soil microbial community that supports enhanced N-cycling functions and plant productivity. Further research is warranted to determine whether similar effects can be achieved in different soil types and climates, which would expand our understanding of the broader applicability of long-term liming and facilitate the prediction of its effects across diverse environments.

## 5. Conclusion

Consistent with our hypothesis, long-term application of lime for 65 years enhanced the abundances of microbial N-cycle genes, including those related to nitrification and denitrification. Furthermore, liming had a buffering effect on the abundances of microbial N-cycle genes, especially for diazotrophs (*nifH)*, significantly mitigating seasonal variations. Importantly, liming did not increase N_2_O emissions, despite the higher abundances of nitrifier (*amoA*) and denitrifier (*nirS* and *nirK*) genes. This absence of increased emissions can be attributed to the enhanced abundance of *nosZ* clade II genes. Our findings suggest that long-term liming in acidic semi-natural grasslands can enhance microbial N-cycling processes and increase plant production without a concomitant increase in N_2_O emissions.

## Acknowledgements

The authors thank Ji Li and Shuaimin Chen for their assistance with field sampling, and Agata S. Pijl for molecular laboratory technical assistance.

